# Hippocampal-midbrain interactions link encoding-related pupil response to memory success

**DOI:** 10.64898/2026.08.10.743973

**Authors:** Alex Kafkas, Hera Y-J Baek, Nanne Kukkonen, Daniela Montaldi

## Abstract

Encoding-related pupil responses predict later memory performance, but the neural mechanisms linking these autonomic dynamics to memory formation remain unclear. This study examined whether pupil responses during encoding track activity in the brain’s memory network and whether they reflect functional interactions between memory-related regions and neural systems involved in pupil control. Participants performed an incidental encoding task involving object stimuli while undergoing simultaneous fMRI and pupillometry; recognition memory was subsequently assessed outside the scanner. Greater pupil constriction during encoding predicted both the strength and quality of later memory. These pupil dynamics correlated with activity in memory-related brain regions, notably the hippocampus and the parahippocampal cortex. Connectivity analyses indicated that encoding-related pupil responses were supported by functional interactions between the hippocampus and the midbrain Edinger–Westphal nucleus, the striatum, and the orbitofrontal cortex. The findings suggest that interactions between memory-related regions and parasympathetic pupil-control systems may modulate encoding efficiency. Together, the results identify encoding-related pupil constriction as a non-invasive marker of memory-network engagement and suggest a hippocampal-midbrain pathway through which autonomic pupil dynamics are coupled with successful memory formation.

## INTRODUCTION

Only some experiences are retained in memory, and understanding why particular events are encoded successfully remains a central goal of cognitive neuroscience. Neuroimaging measures, such as electroencephalography (EEG) and functional magnetic resonance imaging (fMRI), have shown that encoding-related neural processes predict later memory ^1–5^. Pupil responses measured during encoding also reliably predict subsequent memory^6,7^, suggesting that autonomic dynamics may provide a non-invasive window into the physiological state in which memory formation occurs. However, the neural mechanisms linking pupil dynamics to memory formation remain underexplored. The present study therefore used concurrent fMRI and pupillometry to investigate whether encoding-related changes in pupil size are mirrored by activation within the brain’s memory network, offering insight into the physiological pathways that support successful memory formation.

Variations in pupil responses during encoding have been linked to different memory outcomes. Reduced pupil dilation—or constriction—has been found to predict both the type and strength of subsequent memory^6,8–11^. In the first study to demonstrate this link^6^ it was shown that a linear reduction in pupil response, during an incidental encoding task, distinguished forgotten from remembered stimuli, with greater constriction predicting stronger memory. This finding, since replicated^10–12^, aligns with the pupil old/new effect, where reduced pupil dilation is observed for new compared to old stimuli^13–16^. However, opposite patterns have also been reported, particularly under conditions of unexpected novelty, where increased dilation predicted better memory^17–19^. For example, in a recent study^9^, unexpected novel stimuli elicited increased pupil dilation and stronger memory (i.e., recollection), while expected novel stimuli produced constrictions at encoding. These findings suggest that pupil responses reflect the type of novelty encountered, and that task demands shape encoding strategies, associated brain activity, and the resulting pupil dynamics^7,9,15^. The mechanism underlying this remains unclear, however, and specifically, whether the neural system supporting encoding-related pupil changes engages the memory network reflecting communication between pupil control and memory-related brain regions. Addressing this is crucial for understanding the driving mechanisms and evaluating pupillometry as a reliable marker of memory formation.

fMRI studies have demonstrated that subsequent memory is accompanied by activation in the hippocampus and the prefrontal cortex during encoding, particularly for later recall and recollection ^4,22–25^. A hierarchical organisation of the MTL has dominated theories of episodic memory ^26–29^, with the hippocampus supporting associative memory, recall and recollection, and the cortical regions of the MTL, such as the parahippocampal and perirhinal cortices supporting item memory and familiarity-based recognition. Other regions, including parietal, fusiform, and premotor cortices, also predict subsequent memory, though their roles are less well understood. These may provide content-specific (e.g., fusiform) or attentional (e.g., parietal, premotor) support at encoding ^30^, while the MTL, particularly the hippocampus, is central to the formation of new memory representations.

Pupil size is regulated by the autonomic nervous system via two opposing iris muscles: the parasympathetic system, using cholinergic transmission, activates the sphincter muscle to constrict the pupil, while the sympathetic system activates the dilator muscle to induce dilation^31,32^. Sympathetic control originates in the hypothalamus, whereas parasympathetic control is mediated by the Edinger–Westphal (EW) nucleus in the midbrain, which houses preganglionic neurons projecting to the ciliary ganglion^33^. These pathways likely interact with higher cortical structures, allowing cognitive processes to modulate pupil size^34^. The complex modulation of pupil size involves different neurotransmitter systems^35,36^. The parasympathetic system, using acetylcholine, links to cortical and limbic regions and is implicated in learning and synaptic plasticity during memory encoding^37–39^. Cholinergic dysfunction, including in the basal forebrain and possibly the EW, is associated with long-term memory impairment and Alzheimer’s disease^40–42^. The sympathetic system, primarily noradrenergic, is linked to pupil dilation via the locus coeruleus (LC), a key source of noradrenaline and dopamine for the hippocampus and limbic system^43–49^. The LC is also functionally connected to the EW^50^. Taken together, this suggests that midbrain nuclei and associated neurotransmitter activity may mediate the relationship between pupil dynamics and memory encoding processes, which will be investigated more directly here.

We investigated the neural basis of pupil responses associated with successful memory formation. Using an established incidental encoding paradigm with minimal cognitive demand^6^, we examined whether encoding-related changes in pupil size predicted subsequent memory and whether these pupil responses were mirrored by activity in memory-related brain regions. If pupil responses are directly linked to memory formation, activity in regions such as the hippocampus should covary with these responses, and functional connectivity should be observed between the memory network and midbrain structures involved in pupil control.

Alternatively, if encoding-related pupil changes are supported primarily by regions outside the medial temporal memory network, such as frontoparietal or sensory-association regions involved in attention and task-related stimulus processing, this would suggest that pupil responses reflect broader cognitive processes engaged during memory encoding rather than memory-network engagement specifically.

## RESULTS

Nineteen right-handed, healthy participants completed an incidental encoding task during concurrent fMRI and eye tracking, followed by a recognition memory test outside the scanner. During encoding, participants viewed 180 object images while performing an oddball detection task to maintain attention. At test, each item was classified as new, familiar (with graded familiarity strength), or recollected (see Figure 1a and Methods).

**Figure 1.**
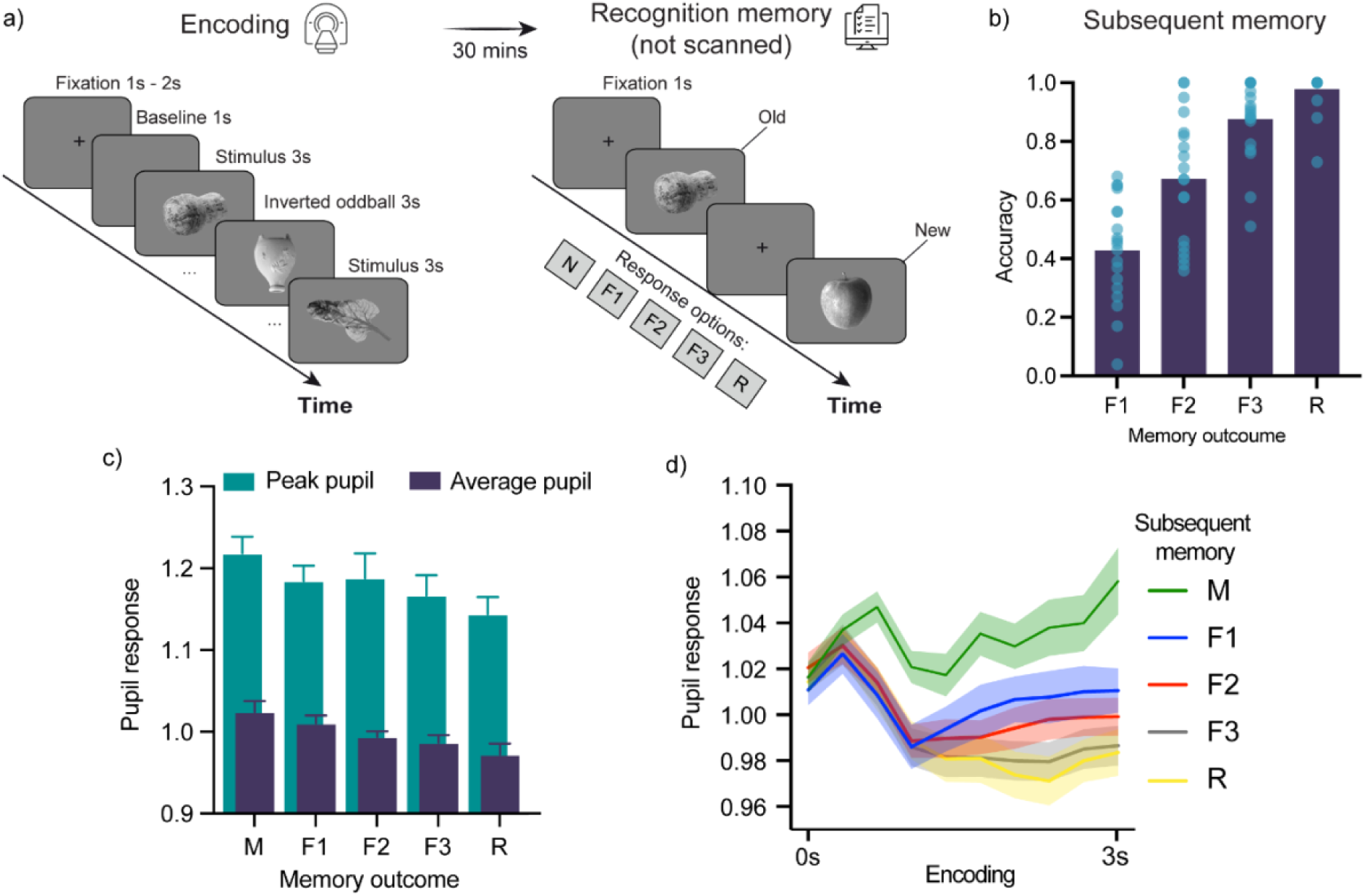
Experimental design, behavioural and pupil effects. a) An incidental encoding task was implemented at encoding while pupil response and fMRI data were collected. A recognition memory task was performed 30 minutes after the end of the encoding task outside the scanner. b) Memory accuracy at recognition across reported memory type. c) Pupil response (peak pupil and average pupil) at encoding predicting different memory outcomes. d) Pupil timeseries during the encoding period for different subsequent memory outcomes. Error bars show the standard error of the mean. M = misses; F1 = weak familiarity; F2 = moderate familiarity; F3 = strong familiarity; R = recollection.

### Behaviouraland pupillometry results

The proportion of responses and response times (RTs) across recognition memory outcomes are summarised in Supplementary Table 1. Memory accuracy increased systematically across familiarity strength (F1, F2, F3; *F*_2,36_ = 50.89, *p* < 0.001, *η*^2^ = 0.74; Figure 1b), and recollection responses were highly accurate (M = 0.98, SD = 0.07). Recollection accuracy exceeded accuracy for F1 (M = 0.43, SD = 0.18; *t*_18_ = 12.22, *p* < 0.001, d’ = 0.20), F2 (M = 0.67, SD = 0.22; *t*_18_ = 6.53, *p* < 0.001, d’ = 0.20), and F3 responses (M = 0.87, SD = 0.13; *t*_18_ = 3.94, *p* < 0.001, d’ = 0.11). RTs also varied with familiarity strength (*F*_2,34_ = 7.34, *p* = 0.002, *η*^2^ = 0.30), with faster F3 responses than both F1 (*p* = 0.002) and F2 responses (*p* = 0.01; Supplementary Table 1). RTs to recollected stimuli did not differ significantly from F1, F2, or F3 responses (all *p*s > 0.05).

Encoding-related pupil responses varied systematically with subsequent memory outcome. Both average and peak pupil responses differed across outcomes (average pupil: *F*_4,68_ = 7.82, *p* < 0.001, *η*^2^ = 0.32; peak pupil: *F*_4,68_ = 3.85, *p* = 0.007). In both measures, this relationship followed a significant linear trend (average pupil: *F*_1,17_ = 12.21, *p* = 0.003; peak pupil: *F*_1,17_ = 14.51, *p* = 0.001): reduced pupil responses predicted stronger familiarity and recollection, whereas missed stimuli showed the largest pupil dilation (Figure 1c). The same pattern was evident in the pupil time-series analysis, which showed a significant memory outcome by time interaction (*F*_36,612_ = 1.86, *p* = 0.002, *η*^2^ = 0.099; Figure 1d).

### fMRI results

#### Subsequent memory effects

Responses to subsequently recognised stimuli engaged a network of regions consistent with prior subsequent-memory findings. Table 1 summarises these effects separately for strongly familiar stimuli (F3 > M) and recollected stimuli (R > M). Within the medial temporal lobe, activity differentiated the two memory outcomes: the left hippocampus showed increased activity for subsequently recollected stimuli, whereas the left parahippocampal cortex showed increased activity for subsequently strongly familiar stimuli.

**Table 1.** Subsequent memory effects in the whole brain for later familiar (F3 > M) and recollected (R > M) stimuli.

| Region | Brodmann area | Voxels | T value | MNI |
| --- | --- | --- | --- | --- |
| **Contrast: $F3 > M$** |  |  |  |  |
| Left fusiform gyrus | BA37/19 | 265 | 7.25 | -39 -67 -16 |
| Right superior parietal gyrus | BA7 | 19 | 6.59 | 33 -58 59 |
| Right superior occipital gyrus | BA19 | 73 | 6.54 | 30 -73 41 |
| Inferior temporal gyrus | BA37 | 36 | 6.48 | 54 -52 -7 |
| Left superior parietal gyrus | BA7 | 16 | 5.6 | -24 -73 44 |
| Inferior occipital gyrus | BA19 | 36 | 5.4 | 39 -79 -4 |
| Left inferior frontal gyrus |  | 14 | 5.18 | -33 29 17 |
| Right inferior frontal gyrus | BA46 | 7 | 4.49<sup>^</sup> | 54 35 17 |
| Left parahippocampal cortex | BA36 | 17 | 4.81 | -30 -34 -22 |
| **Contrast: $R > M$** |  |  |  |  |
| Left fusiform gyrus and middle occipital gyrus | BA37/19 | 407 | 9.5 | -39 -76 -16 |
| Right fusiform gyrus and middle occipital gyrus | BA37/19 | 303 | 8.81 | 51 -49 -10 |
| Right superior parietal gyrus | BA7 | 108 | 7.95 | 27 -67 50 |
| Left inferior frontal gyrus | BA9 | 28 | 6.69 | -45 5 32 |
| Left inferior parietal gyrus | BA7 | 22 | 6.43 | -27 -55 53 |
| Right inferior parietal gyrus | BA40 | 6 | 4.8<sup>^</sup> | 42 -34 32 |
| Right inferior frontal gyrus | BA46 | 48 | 6.02 | 48 26 17 |
| Right supramarginal gyrus | BA40 | 42 | 5.77 | 48 -34 44 |
| Left inferior occipital gyrus | BA18 | 5 | 5.46<sup>^</sup> | -24 -88 -19 |
| Right calcarine sulcus | BA17 | 30 | 5.38 | 12 -97 -1 |
| Right inferior frontal gyrus |  | 20 | 5.28 | 51 8 26 |
| Right middle occipital gyrus | BA19 | 23 | 4.54 | 33 -76 20 |
| Left middle occipital gyrus | BA19 | 13 | 5.00 | -30 -67 23 |
| **Left supramarginal gyrus** | BA40 | 7 | 4.52<sup>^</sup> | -60 -28 44 |
| **Left hippocampus (posterior)** |  | 6 | 4.04<sup>^</sup> | -30 -34 -1 |
| **Left hippocampus (anterior)** |  | 10 | 4.42 | -15 -10 -22 |

*Note:* Activations are significant at a cluster-corrected FWE of $p < 0.05$ using nonparametric permutations, with the exception of those marked with <sup>^</sup> which are significant at an uncorrected threshold of $p < 0.001$ .

#### Brainactivity modulatedby pupil response

The pupil response patterns reported above replicate previously reported effects using the same incidental encoding task^6^. Moreover, the activation patterns supporting recollection and familiarity implicate regions consistent with the established neural bases of these kinds of memory, including the hippocampus and parahippocampal cortex, respectively. To investigate the neural basis of encoding-related pupil changes and their overlap with memory networks, parametric analyses were conducted across all trial types, regardless of memory outcome.

Separate parametric models also examined activity as a function of pupil response for all hits, recollections, and strongly familiar stimuli. Because decreased pupil responses predicted subsequent memory strength and type and because the present study sought to identify the neural basis of this established behavioural effect, the primary analyses focused on negative modulation by pupil response, that is, increased activity associated with reduced pupil response. Activations related to the positive modulation by pupil response, reflecting increased activity with increased pupil response, are reported in the Supplement (Suppl. Table 4) but do not change the proposed interpretation of our findings.

Parametric modulation by decreased pupil response for all hits revealed activations in the bilateral hippocampus (MNI: -24 -16 -19 and MNI: 30 -13 -22, both *p*_FWE_ < 0.05), right superior frontal lobe (MNI: 6 -22 65, *p*_FWE_ < 0.05), bilateral lingual gyrus (MNI: 21 -88 -1, *p*_FWE_ < 0.05; MNI: -30 -91 -7, *p*_uncorrected_ < 0.001), and right fusiform gyrus (MNI: 27 -61 -10, *p*_uncorrected_ < 0.001; see Suppl. Table 2). Activity in these regions therefore tracked decreases in pupil response associated with later hits, with the strongest inference supported by the FWE-corrected hippocampal, frontal, and lingual effects. A comparable analysis across all trials, including hits and misses, revealed similar bilateral hippocampal activations (MNI: -30 -13 -22 and MNI: 30 -13 -22, both *p*_FWE_ < 0.05), right superior frontal lobe activations (MNI: 36 -22 50 and MNI: 6 -22 59, both *p*_FWE_ < 0.05), and lingual (MNI: -24 -88 -13, *p*_FWE_ < 0.05) and fusiform gyrus activity (MNI: 30 -49 -16, *p*_uncorrected_ < 0.001). This analysis also identified the left parahippocampal cortex (MNI: -24 -34 -22, *p*_FWE_ < 0.05). When recollected trials were analysed separately, the right hippocampus (MNI: 24 -13 -19, *p*_FWE_ < 0.05) tracked decreases in pupil response. By contrast, strongly familiar trials involved activity in the parahippocampal cortex (MNI: -18 -40 -10, *p*_FWE_ < 0.05) and amygdala (MNI: -12 -4 -22, *p*_uncorrected_ < 0.001) as a function of pupil response change (Figure 2 and Suppl. Table 2). These findings indicate that distinct medial temporal lobe regions tracked encoding-related pupil responses depending on the form of later memory, with hippocampal effects most evident for recollection and parahippocampal effects most evident for strong familiarity.

**Figure 2.**
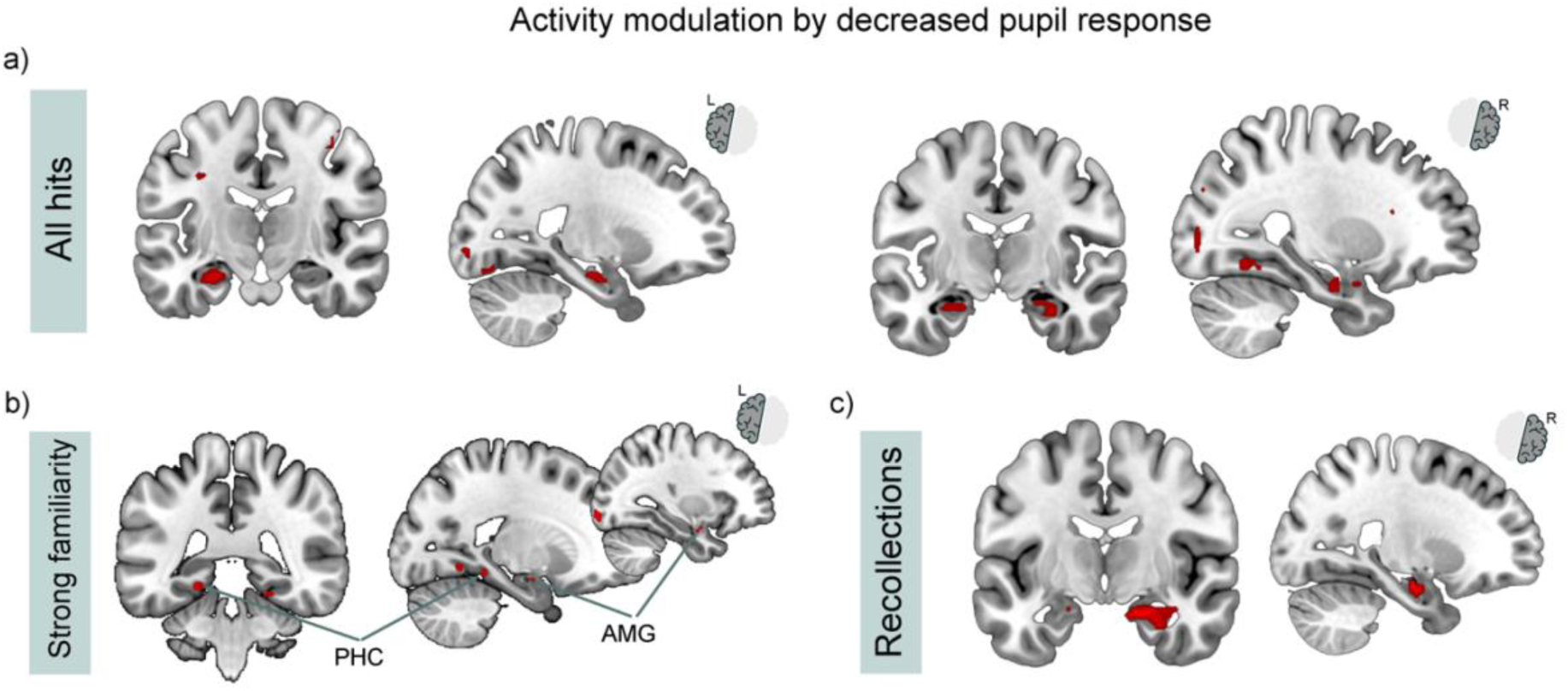
**Hippocampal (a and c) and other** medial temporal lobe activity (b) associated with decreased pupil response (constriction) at encoding for different memory outcomes. a) All hits, b) **strongly** familiar stimuli, and c) **recollections.** Activations are thresholded at *p*_FWE_ < 0.05, unless otherwise stated in the text and Suppl. Table 2. PHC = parahippocampal cortex; AMG = amygdala.

#### Psychophysiological interaction with hippocampal seeds

Next, we explored functional connectivity from the hippocampal regions that showed significant activity as a function of pupil response for all hits and for recollections alone. A PPI analysis using the left hippocampus (-24 -16 -19) as a seed, with parametric modulation as a function of pupil response for hits, revealed connectivity with the right hippocampus (MNI: 30 - 22 -10, *p*_FWE_ < 0.05) and a cluster within the left superior temporal gyrus (MNI: -39 -55 14, *p*_FWE_ < 0.05). The corresponding analysis using the right hippocampal seed (30 -13 -22) revealed increased connectivity with the left striatum (MNI: -18 -10 -4, *p*_FWE_ < 0.05), left inferior orbitofrontal gyrus (MNI: -36 23 23 and MNI: -42 23 -4, both *p*_uncorrected_ < 0.001), left insula (MNI: -27 29 -1, *p*_FWE_ < 0.05), left superior frontal gyrus (MNI: -18 2 65, *p*_FWE_ < 0.05), and a right midbrain region (MNI: 3 -16 -16, *p*_FWE_ < 0.05) encompassing the vicinity of the red nucleus and Edinger-Westphal region (Suppl. Table 3 and Figure 3a).

**Figure 3.**
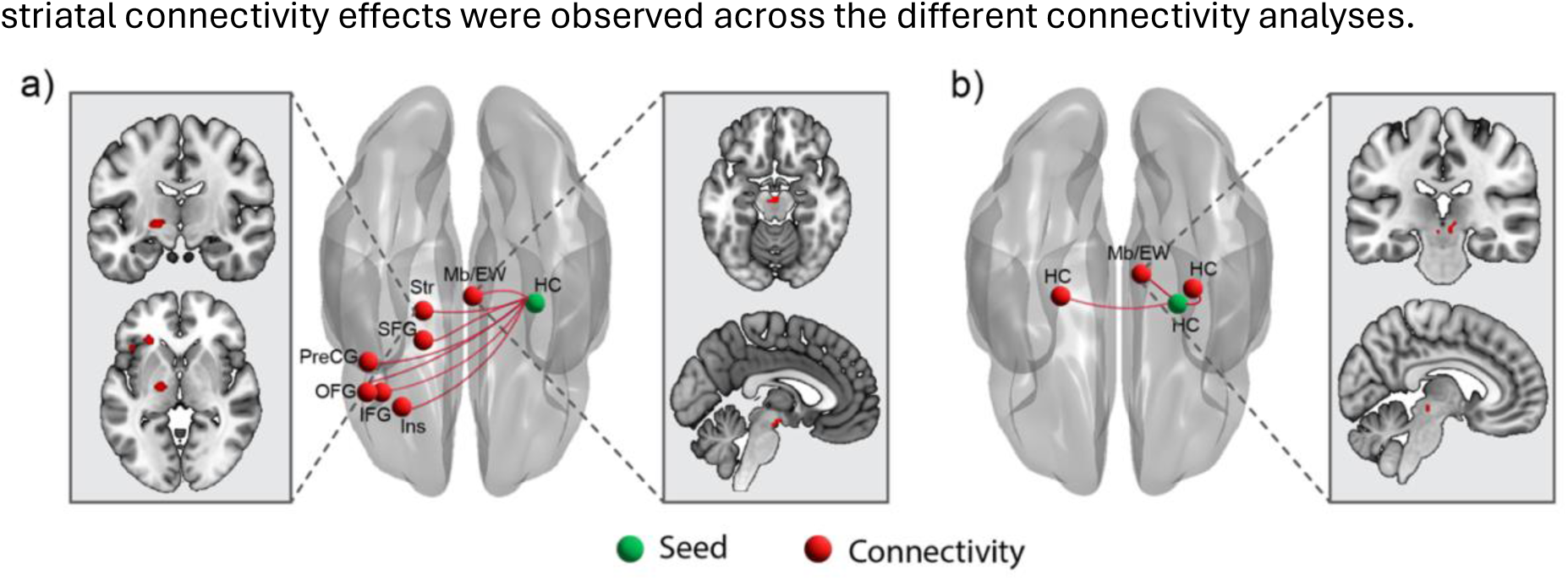
Significant connectivity modulation effects associated with encoding-related pupil constrictions from the right hippocampus to other brain regions. a) Connectivity patterns from the right hippocampus (30 -13 -22) for hits and b) **connectivity** patterns from the right hippocampus (24 -13 -19) for recollected stimuli. HC = hippocampus; Mb/EW = midbrain and Edinger-Westphal nucleus; Str = striatum; SFG = superior frontal gyrus; PreCG = precentral gyrus; OFG = orbitofrontal cortex; IFG = inferior frontal gyrus; Ins = insula. Activations are thresholded at *p*_FWE_ < 0.05, unless otherwise stated in the text. All MNI coordinates and statistical details related to these activations are presented in Supplementary Table 3.

The connectivity analyses restricted to recollections, using the right hippocampal seed (24 -13 -19), revealed connectivity with the bilateral hippocampus (MNI: 30 -19 -7, *p*_FWE_ < 0.05; MNI: -24 -16 -13, *p*_uncorrected_ < 0.001) and bilateral midbrain regions (MNI: 9 -25 -4, *p*_FWE_ < 0.05; MNI: -3 -25 -10, *p*_uncorrected_ < 0.001) in the vicinity of the red nucleus and Edinger-Westphal region (Suppl. Table 3; Figure 3b). As expected, a similar but slightly more extensive connectivity network was observed when the analysis was conducted across all stimulus types, irrespective of memory outcome; these results are presented in Supplementary Table 3. Taken together, the PPI analyses suggest that hippocampal interactions with midbrain and striatal regions covaried with encoding-related pupil response changes, rather than establishing the direction of influence between these regions. Critically, hippocampal-to-hippocampal, midbrain, and striatal connectivity effects were observed across the different connectivity analyses.

## DISCUSSION

The present study investigated the neural substrates of encoding-related pupil responses associated with the experience of successful memory. The findings show that reduced pupil dilation, or pupil constriction, predicted both subsequent memory strength and memory type. This study replicates previous findings using a similar incidental encoding paradigm^6^ and extends them by showing that encoding-related pupil responses were associated with activity in the memory network, particularly the hippocampus and parahippocampal cortex.

Connectivity analyses further indicated that pupil-linked memory effects covaried with functional interactions between the hippocampus and prefrontal, striatal, and midbrain regions, including the Edinger–Westphal region. Together, these findings suggest that encoding-related pupil constriction reflects a set of coordinated interactions between medial temporal lobe and midbrain systems during successful memory formation.

Within the MTL, hippocampal activation at encoding predicted recollection, while parahippocampal cortex activation predicted familiarity, mirroring the functional specialisation established at retrieval^14,51^. The present results extend this dissociation by showing that pupil constriction was associated with increased hippocampal activity, particularly for subsequently recollected stimuli, whereas pupil changes linked to subsequent familiarity involved the parahippocampal cortex and amygdala^27^. These findings suggest that encoding-related pupil responses are sensitive to the balance of hippocampal and parahippocampal engagement during successful memory formation, with stronger hippocampal involvement when associative encoding supports later recollection.

Our findings further identify a candidate pathway linking parasympathetic pupil-control systems with memory formation. Encoding-related pupil constriction was associated with stronger functional connectivity between the hippocampus and a midbrain region consistent with the EW region, a key component of parasympathetic pupil control. The EW, particularly its preganglionic cholinergic neurons^52^, controls pupil constriction via input from the locus coeruleus^50,53,54^. Recent evidence shows that ventral hippocampal neurons can directly influence EW activity to support attention^55^. Although the present PPI analyses cannot establish the direction of influence between the hippocampus and the EW region, this anatomical evidence provides one plausible route by which hippocampal engagement during memory encoding may interact with parasympathetic pupil-control systems. Conversely, EW-mediated pupil dynamics may index the autonomic state in which hippocampal encoding occurs. Thus, the present findings are best interpreted as evidence for coordinated hippocampal–midbrain interactions that link memory-network engagement with parasympathetic pupil control during successful encoding. Pupil constriction patterns may therefore provide sensitive, real-time markers of memory-network activity and reflect the strength and quality of incidental encoding.

The present study purposefully used an incidental encoding task paired with a simple detection task that placed minimal cognitive demands on participants. This design was well suited to detecting encoding-related pupil responses because it minimised additional decision-related or strategic encoding demands. By contrast, intentional encoding paradigms or more demanding decision-making tasks may obscure pupil-based subsequent memory effects by introducing pupil changes related to effort, decision complexity, or task strategy. Consistent with this possibility, studies using complex encoding tasks or manipulating multiple dimensions have often failed to detect pupil-based subsequent memory effects (see e.g.^16,56^).

Of particular relevance to the present study is the theoretical distinction between stimulus novelty – novel items within an expected context – and contextual novelty – unexpected combinations of familiar and novel elements^15,21^. Previous evidence^9^ indicates that novelty expectancy, that is, whether a new stimulus is expected or unexpected, reliably modulates pupil patterns that predict subsequent memory, suggesting that different encoding processes are triggered depending on the type of novelty encountered. Consistent with earlier work^6^, we propose that when expected novelty is detected, cholinergic interactions between the memory network and the parasympathetic system are engaged to support successful memory formation. The present findings support this account, particularly the observed connectivity between the hippocampus and EW region. Future research should extend this work by investigating encoding-related pupil dynamics under conditions of unexpected novelty and surprise. Building on previous behavioural^9,21^ and fMRI findings^18^, we hypothesise that unexpected novelty will engage distinct hippocampal–midbrain connectivity patterns, reflecting different neurotransmission profiles and autonomic activity compared to those triggered by the current task.

The exact mechanisms by which novelty is assessed upon encountering new information remain unclear. Evidence consistently highlights the hippocampus as central to the evaluation and encoding of novelty^15,18,57–59^. The current study’s connectivity analyses are consistent with this view, showing hippocampal-to-hippocampal connectivity involving anterior, midline/posterior, and contralateral hippocampal regions. This connectivity pattern may reflect the neural computations involved in evaluating the nature of novelty and the subsequent encoding of relevant new events^60^. Alternatively, it may reflect a two-stage mechanism in which initial novelty signals in the hippocampus propagate to other brain regions before integrated outcomes feed back to the hippocampus^57^.

Although the hippocampus appears central to novelty evaluation, this process likely depends on interactions with a broader network. Here, we identified potential component regions, notably the striatum and orbitofrontal cortex (OFC). The OFC has previously been shown to interact with the hippocampus during olfactory sequence learning in rats^61,62^ and maintains functional connectivity with the human hippocampus via cholinergic inputs from the nucleus basalis^63^. Similarly, the striatum’s role in memory formation involves functional interplay with the hippocampus^18,64,65^. A recent rodent study further supports cholinergic engagement of both hippocampal and striatal areas, including the globus pallidus and putamen^66^, during memory formation. Consistent with this broader literature, the current results demonstrate that hippocampal-striatal connectivity covaried with encoding-related pupil response patterns that predicted successful memory. Collectively, the findings point to a coordinated network encompassing hippocampal, parahippocampal, striatal, midbrain, and orbitofrontal regions supporting pupil constriction and the neural activity associated with novelty and memory, although the neurotransmitter mechanisms require further exploration.

An important consideration is whether the identified pathway is exclusively cholinergic. Although the present findings cannot definitively answer this, substantial evidence supports the cholinergic hypothesis. Observed pupil constriction aligns with parasympathetic, predominantly cholinergic control of the eye’s constrictor muscle^33,53^, with the EW nucleus identified as a key cholinergic source^52^. Other connected regions, such as the orbital and inferior prefrontal cortex^63^ and the striatum^66^, receive cholinergic inputs vital for hippocampal-dependent learning. Nonetheless, the influence of other neurotransmitters requires further clarification. For instance, GABAergic activity from the globus pallidus^67^ might inhibit dopaminergic or noradrenergic pathways to the hippocampus. Supporting this, optogenetic studies in mice indicate that GABAergic neurons in the locus coeruleus suppress noradrenergic-based pupil dilation^68^. Future human studies, potentially combining high-resolution MR spectroscopy with pupillometry, are warranted to examine these interactions further.

The modest sample size should also be considered when interpreting the connectivity findings. Although the effects were detected using non-parametric cluster correction and converged across behavioural, pupillometric, activation, and connectivity analyses, the PPI results should be treated as identifying a candidate hippocampal–midbrain pathway that requires replication in larger samples. In addition, PPI analyses cannot establish the directionality or causal nature of the observed interactions. Given the small size of the EW nucleus and its proximity to neighbouring midbrain structures, these effects should be interpreted as implicating the EW region rather than selectively localising the nucleus itself. Nevertheless, the observation of convergent effects despite the modest sample size provides some reassurance about the robustness of the reported associations.

Taken together, our results replicate the finding that pupil responses can serve as reliable predictors of the type and strength of subsequent memory, while also clarifying the neural basis of this relationship. By showing that pupil constriction patterns are associated with activity in the hippocampus and broader memory network, we highlight the role of autonomic nervous system interactions in memory formation. Moreover, functional connectivity between the hippocampus and the Edinger–Westphal region identifies a candidate pathway through which memory-network activity and parasympathetic pupil-control systems may be coupled during encoding. This demonstrates the value of pupillometry in the detailed exploration of the neural bases of memory. Future pupillometry studies are needed to test the causal and neurotransmitter-specific mechanisms underlying this pathway and to explore potential applications in healthy ageing and memory disorders.

## METHODS

### Participants

Twenty healthy participants were recruited from the University of Manchester and the Greater Manchester area via campus adverts and the University’s online volunteer recruitment platform. All participants provided written informed consent prior to participation. Data from one participant were excluded because an experimental failure during scanning resulted in partial scan completion only, leaving 19 participants for analysis (12 females; mean age = 25.7 years, SD = 3.60). All participants were right-handed, native English speakers, had normal or corrected-to-normal vision with contact lenses, and self-reported no history of neurological or psychiatric conditions. None was taking psychotropic medication or had a history of drug addiction, and all were asked to abstain from alcohol for 24 hours before participation.

Participants were compensated at a rate of £20 per session. All procedures were approved by the University of Manchester Research Ethics Committee.

### Stimulus material

The stimulus material consisted of 300 object pictures (500 × 375 pixels) from Kafkas and Montaldi^6^. The stimulus set comprised greyscale pictures of single everyday objects depicting non-emotional, man-made and natural items, such as a stapler or onion. The set was matched for low-level visual properties, including luminosity, luminance, and chromatic information, making it appropriate for pupillometry studies^6,69^. At encoding, 180 stimuli were presented, comprising 90 man-made and 90 natural objects, along with 20 inverted oddballs (see Procedure). At retrieval, 120 studied stimuli were presented with 90 new stimuli, yielding 210 test trials in total. Ten additional stimuli were used for practice before the encoding and retrieval blocks.

### Procedures

The study consisted of an encoding task completed in the MRI scanner and a recognition memory task completed after the scanning session (see Figure 1). An incidental encoding task was used to replicate the procedure developed by Kafkas and Montaldi^6^. During scanning, participants were instructed to look carefully at each stimulus and press a button whenever an inverted item appeared. This oddball detection task was used to maintain attention during encoding. Data from the 20 oddball stimuli were excluded from further analysis, and all analyses reported here concern the 180 incidentally encoded object stimuli. Concurrent eye-tracking data were recorded using an EyeLink 1000 Plus remote eye tracker. Each trial began with a central fixation cross lasting 1–2 s, followed by a grey baseline screen for 1 s, which was used to calculate trial-specific pupil responses (see Eye-tracking recording and preprocessing). Each object was then presented for 3 s, during which participants could indicate whether it was inverted by pressing a button. The next trial began after a fixation cross with a mean intertrial interval of 1.5 s (range: 1–2 s), optimised for EPI acquisition. Null events consisting of an implicit crosshair baseline lasting 3 s were intermixed with object trials to optimise jittering and provide baseline BOLD measures.

Approximately 30 minutes after the end of the encoding task, participants completed a recognition memory task in a testing room outside the scanner. Before the task, participants were trained to distinguish studied from unstudied stimuli and to discriminate familiarity from recollection for studied items. They were instructed to classify a stimulus as familiar when they recognised it as having been presented at encoding, and as recollected when they could retrieve associative information from the encoding episode (for detailed instructions, see ref^27^). Participants practised these decisions and could ask questions before the main task began.

During the recognition memory task, each previously studied or unstudied stimulus was presented for 3 s and classified as familiar, recollected, or new. For familiar responses, participants rated familiarity strength as weak (F1), moderate (F2), or strong (F3). The task comprised 210 stimuli presented one at a time in random order, including 120 old and 90 new stimuli. Responses were made on a keyboard using five buttons corresponding to new, F1, F2, F3, and recollection.

### fMRI data acquisition and preprocessing

MRI data were collected using a 3T Philips Achieva scanner. Functional images were acquired using gradient echo-planar imaging with blood oxygenation level-dependent contrast. A total of 468 volumes were recorded per participant across three sessions, with each volume comprising 40 slices (TR = 2.5 s; TE = 35 ms; matrix size = 80 × 80; voxel size = 3 × 3 × 3.5 mm) covering the whole brain. High-resolution T1-weighted images were obtained at the beginning of each session before the functional blocks, comprising 144 slices with an isotropic voxel size of 1 mm and a matrix size of 256 × 256.

The quality of the EPI time-series data was assessed using ArtRepair. Residual movement parameters were included in the GLM models. Preprocessing was performed in SPM12 (Statistical Parametric Mapping, Wellcome Trust Centre for Neuroimaging). Functional EPI data were realigned to the mean image using a six-parameter rigid-body transformation and resliced using sinc interpolation to reduce motion-induced artefacts in the fMRI time series. Slice-timing correction was applied to the resliced images to correct for differences in acquisition time across slices. Each participant’s high-resolution T1 image was coregistered to the corresponding mean EPI image. After spatial normalisation, the functional data were resliced to 3 mm isotropic voxels and spatially smoothed using a 6 mm isotropic full-width at half-maximum Gaussian kernel.

### fMRI data analyses

Encoding trials were sorted by subsequent memory outcome (misses and hits: F1, F2, F3, and R) separately for each participant. Event-related BOLD responses were modelled using a canonical haemodynamic response function convolved with the onset times and durations of each event or trial type^70^. Three first-level GLMs were estimated to examine brain responses associated with pupil changes during encoding. The first GLM included all encoded trials, with pupil response specified as a parametric modulator convolved with each event’s haemodynamic response function^71^. The second GLM modelled hit events (accurate F1, F2, F3, and R responses) and misses as separate conditions, each with pupil response as a parametric modulator. The third GLM modelled F1, F2, F3, R, and miss trials as separate conditions, with pupil response included as a parametric modulator for each condition. In all models, nuisance regressors included trials with no valid pupil response (<5% of trials), inverted oddball trials, six movement parameters from realignment for each functional run, and residual movement artefacts identified by ArtRepair. Low-frequency noise was removed using a 128 s high-pass filter.

To identify brain regions responding to encoding-related pupil changes, first-level parametric *t*-contrasts were created to test monotonic increases in activity associated with decreased pupil response, that is, pupil constriction. Decreased pupil responses were prioritised because they predicted subsequent memory in both the present study and previous research using the same paradigm^6^. Activations associated with increased pupil response during encoding were also analysed and are reported in the Supplement (Table 4). Nonlinear quadratic effects were modelled to capture residual variance not explained by the linear parametric contrasts, but they did not produce significant additional activations and are not reported separately. Subsequent-memory effects associated with recollection and strong familiarity were examined using two directional *t*-contrasts: R > M and F3 > M.

In the second-level analysis, individual contrasts were treated as random effects and combined at the group level. One-sample *t*-tests were performed with an initial voxel-wise threshold of *p* < 0.001, and cluster-level family-wise error correction was set at *p*_FWE_ < 0.05 using non-parametric permutation testing with 5,000 permutations in SnPM13. In selected cases, uncorrected *p* < 0.001 effects are reported to describe subthreshold activity in small brain regions, such as midbrain nuclei. These instances are explicitly noted in the text and relevant tables.

### Psycho-physiological Interaction (PPI) analysis

PPI analysis^72^ was performed to identify regions whose functional connectivity with hippocampal seed regions covaried with decreased pupil responses during successful memory formation. The main GLM parametric modulation analyses showed bilateral hippocampal activation associated with decreased pupil responses, that is, negative modulation by pupil response. The hippocampus was therefore selected as a functional seed region for the PPI analyses. The left hippocampus (x = -24, y = -16, z = -19) and right hippocampus (x = 30, y = -13, z = -22) were used as seed regions in separate PPI analyses for hit responses. A separate recollection-specific PPI analysis was also conducted using the right hippocampal seed region (x = 24, y = -13, z = -19). Deconvolved BOLD activity was extracted from these regions of interest for the pupil-response parametric modulation models described above.

For each seed region, first-level PPI GLMs were constructed with three regressors of interest: the seed-region BOLD time series as the physiological regressor, the pupil parametric modulation contrast as the psychological regressor, and their interaction term as the PPI regressor. After model estimation, *t*-contrasts of the interaction term were generated for each participant and analysed at the group level using a random-effects model. Activations from these contrasts, indicating regions whose connectivity with hippocampal seeds was modulated by pupil response, were initially thresholded at *p* < 0.001, and statistical significance was determined using cluster-level family-wise error correction at *p*_FWE_ < 0.05 via non-parametric permutation testing in SnPM13, unless otherwise stated.

### Eye-tracking recording and preprocessing

Concurrent with fMRI, eye-tracking data were recorded using an MR-compatible long-range EyeLink 1000 Plus eye tracker (SR Research, Ontario, Canada) at a sampling rate of 1000 Hz. Before data collection, a standard 9-point calibration was performed for each participant. Trial timestamps marked the onset of each baseline period and stimulus presentation. Pupil traces were classified as valid traces, blinks, or partial eyelid closures; blinks and partial eyelid closures were discarded. For each trial, the grand mean pupil trace was calculated, and pupil recordings deviating by more than three standard deviations from this mean were identified as artefacts and discarded. These discarded traces were primarily due to partial eyelid closures occurring immediately before or after blinks. Trials with more than 50% discarded samples were excluded from analysis. Participants with more than 40% excluded trials were also removed from the study. Because discarded recordings were minimal in the final participant sample, interpolation was not applied.

Peak pupil response was calculated by averaging the three pupil samples before and the three samples after each trial’s maximum pupil value, reducing reliance on a single sample and making the measure less vulnerable to noise. Average pupil response was calculated as the mean pupil size across the full trial period. For each trial, both peak and average pupil values were baseline-corrected using the mean pupil diameter during the 1,000 ms grey-screen baseline preceding stimulus onset. Thus, the pupil response to each encoding stimulus was quantified as either the maximal deviation from baseline (peak pupil response) or the mean deviation from baseline (average pupil response).

### Behavioural and pupillometry data analyses

Memory accuracy [hit rate / (hit rate + false-alarm rate)] was calculated for each participant and analysed using a one-way ANOVA with familiarity strength (F1, F2, F3) as a within-subject factor. Recollection accuracy was compared with familiarity accuracy at each strength level using separate *t*-tests. The same analyses were applied to response times. Peak and average pupil responses were analysed using separate one-way ANOVAs with memory outcome (miss, F1, F2, F3, R) as the within-subject factor. The significance level for all statistical comparisons was set at α = 0.05.

## STATEMENTS

### Funding

This study was supported by an Imaging Facilities Grant from the University of Manchester to AK and DM (Ref: 18/07) and by a Wellcome Trust Grant (094597/Z/10/Z) awarded to DM.

### Competing interests

The authors declare no competing interests.

### Data availability

Data supporting the findings of this study are available from the corresponding author upon request.

## SUPPLEMENTARY MATERIAL

**Supplementary Table 1.**
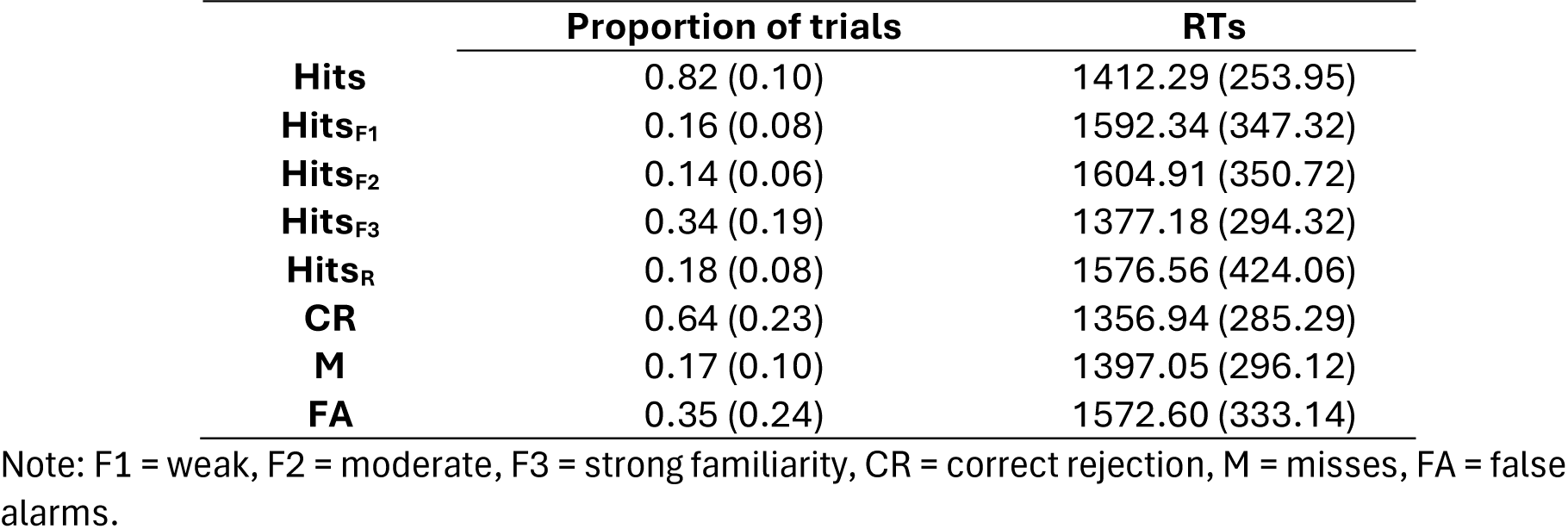
Proportion of trials and response times (RTs) across recognition memory outcomes

**Supplementary Table 2.** Parametric (monotonic) increases in activity at encoding as a function of decreased pupil response across different memory outcomes and for all trials irrespective of memory outcome

| Region | Brodmann area | Voxels | T-value | MNI |
| --- | --- | --- | --- | --- |
| **Hits** |  |  |  |  |
| Right medial frontal gyrus | BA6 | 113 | 5.07 | 6 -22 65 |
| Right somatosensory cortex | BA3 | 101 | 4.51 | 39 -28 53 |
| Left hippocampus |  | 49 | 4.72 | -24 -16 -19* |
| Right hippocampus |  | 17 | 3.59 | 30 -13 -22* |
| Right lingual gyrus | BA17/18 | 44 | 3.89 | 21 -88 -1 |
| Left lingual gyrus | BA17/18 | 29 | 3.56^ | -30 -91 -7 |
| Right fusiform | BA37 | 19 | 3.46^ | 27 -61 -10 |
| **Recollections** |  |  |  |  |
| Right hippocampus |  | 61 | 5.15 | 24 -13 -19* |
| Left hippocampus |  | 5 | 3.17^ | -12 -4 -22 |
| **F3** |  |  |  |  |
| Right lingual gyrus | BA18 | 12 | 3.74 | 24 -88 -1 |
| Left lingual gyrus | BA18 | 11 | 3.70 | -30 -91 -7 |
| Left parahippocampal cortex | BA36 | 8 | 3.74 | -18 -40 -10 |
| Left amygdala |  | 7 | 3.30^ | -30 -4 -22 |
| **Across all trials** |  |  |  |  |
| Right superior frontal gyrus | BA6/7 | 151 | 5.29 | 36 -22 50 |
| Right superior medial frontal gyrus | BA6 | 134 | 5.09 | 6 -22 59 |
| Left hippocampus |  | 60 | 4.69 | -30 -13 -22 |
| Right hippocampus |  | 36 | 4.44 | 30 -13 -22 |
| Left lingual gyrus | BA18 | 76 | 3.77 | -24 -88 -13 |
| Right superior occipital gyrus | BA19 | 30 | 3.76 | 27 -82 29 |
| Left parahippocampal cortex | BA36 | 11 | 3.68 | -24 -34 -22 |
| Left superior occipital gyrus | BA7/19 | 62 | 3.63^ | -18 -82 32 |
| Right fusiform gyrus | BA37 | 35 | 3.57<sup>^</sup> | 30 -49 -16 |
| Right middle occipital gyrus | BA17/18 | 24 | 3.51<sup>^</sup> | 24 -88 5 |

*Note:* Activations are significant at a cluster-corrected FWE of $p < 0.05$ using nonparametric permutations, with the exception of those marked with <sup>^</sup> which are significant at an uncorrected threshold of $p < 0.001$ . \*hippocampal regions used as seeds in the subsequent PPI analyses.

**Supplementary Table 3.** Regions showing increased connectivity with the hippocampus under the influence of encoding-linked pupil response reductions for hits, recollections and collapsed across all trials

| Region | Brodmann area | Voxels | T-value | MNI |
| --- | --- | --- | --- | --- |
| ***Seed left hippocampus [-24 -16 -19] - hits*** |  |  |  |  |
| Right hippocampus |  | 5 | 3.68 | 30 -22 -10 |
| Left superior temporal gyrus | BA22 | 5 | 3.70 | -39 -55 14 |
| ***Seed right hippocampus [30 -13 -22] - hits*** |  |  |  |  |
| Left superior frontal gyrus | BA6 | 24 | 4.64 | -18 2 65 |
| Left insula |  | 20 | 4.56 | -27 29 -1 |
| Left precentral gyrus | BA9 | 11 | 3.92 | -42 11 29 |
| Left striatum (lentiform nucleus and medial globus pallidus) |  | 17 | 3.85 | -18 -10 -4 |
| Right midbrain (Edinger–Westphal nucleus) |  | 8 | 3.76 | 3 -16 -16 |
| Left inferior frontal gyrus | BA46 | 11 | 3.4<sup>^</sup> | -36 23 23 |
| Left orbitofrontal gyrus |  | 5 | 3.45<sup>^</sup> | -42 23 -4 |
| ***Seed right hippocampus [24 -13 -19] – recollections*** |  |  |  |  |
| Right hippocampus |  | 5 | 4.63 | 30 -19 -7 |
| Right midbrain (red nucleus/Edinger–Westphal nucleus) |  | 5 | 3.68 | 9 -25 -4 |
| Left hippocampus |  | 5 | 3.48<sup>^</sup> | -24 -16 -13 |
| Left midbrain |  | 2 | 3.3<sup>^</sup> | -3 -25 -10 |
| ***Seed left hippocampus [-24 -16 -19] – all trials*** |  |  |  |  |
| Right superior parietal lobe | BA7 | 36 | 4.81 | 21 -61 59 |
| Right precentral gyrus | BA4 | 22 | 4.45 | 36 -22 53 |
| Right hippocampus |  | 17 | 4.23 | 30 -25 -16 |
| Left hippocampus |  | 7 | 4.08 | -27 -25 -16 |
| Right postcentral gyrus | BA2 | 24 | 4.05 | 39 -40 62 |
| Left superior frontal gyrus | BA2 | 12 | 3.75 | -15 2 56 |
| Right insula |  | 9 | 3.71 | 48 8 -4 |
| Left middle frontal gyrus | BA10 | 6 | 3.67<sup>^</sup> | -42 44 14 |
| Left lateral globus pallidus |  | 2 | 3.57<sup>^</sup> | -24 -7 -4 |
| Left amygdala |  | 3 | 3.53<sup>^</sup> | -24 -7 -16 |
| Left postcentral gyrus |  | 7 | 3.59<sup>^</sup> | -12 -37 77 |
| Right midbrain |  | 5 | 3.47<sup>^</sup> | 15 -13 -22 |
| ***Seed right hippocampus [30 -13 -22] – all trials*** |  |  |  |  |
| Right striatum (medial GP and putamen) |  | 32 | 5.5 | 15 -13 -4 |
| Left striatum (medial GP and putamen) |  | 24 | 4.03 | -18 -16 -4 |
| Left orbitofrontal gyrus | BA47 | 53 | 4.59 | -27 32 -4 |
| Left superior frontal gyrus | BA6/2 | 28 | 4.21 | -18 5 65 |
| Right fusiform gyrus | BA37 | 8 | 3.87 | 27 -49 -19 |
| Left midbrain (red nucleus/Edinger–Westphal nucleus) |  | 13 | 3.74 | -3 -19 -13 |
| Right insula |  | 17 | 3.69 | 36 8 -4 |
| Left insula |  | 7 | 3.55^ | -36 8 -7 |

*Note:* Activations are significant at a cluster-corrected FWE of $p < 0.05$ using nonparametric permutations, with the exception of those marked with ^ which are significant at an uncorrected $p < 0.001$ .

**Supplementary Table 4.** Parametric (monotonic) increases in activity at encoding as a function of increased pupil response across different memory outcomes and for all trials irrespective of memory outcome

| **Region** | **Brodmann area** | **Voxels** | **T-value** | **MNI** |
| --- | --- | --- | --- | --- |
| ***Hits*** |  |  |  |  |
| Right anterior cingulate | BA32 | 343 | 6.08 | 3 41 14 |
| Anterior thalamus |  | 24 | 5.95 | -9 -13 11 |
| Right angular gyrus | BA39 | 133 | 5.54 | 57 -46 32 |
| Left precuneus | BA7 | 64 | 5.39 | -6 -67 47 |
| Right insula |  | 24 | 5.13 | 36 8 -16 |
| Right superior medial frontal gyrus | BA8 | 28 | 5.1 | 12 38 44 |
| ***Recollections*** |  |  |  |  |
| Right middle cingulate gyrus | BA32/33 | 38 | 5.4 | 9 17 38 |
| Left inferior frontal gyrus | BA45 | 16 | 5.00 | -51 2 23 |
| ***F3*** |  |  |  |  |
| Right anterior cingulate | BA33 | 101 | 6.58 | 6 35 29 |
| Left superior frontal gyrus | BA6 | 64 | 6.87 | -6 11 68 |
| Right frontal pole | BA10 | 34 | 5.31 | 39 47 20 |
| Left caudate and thalamus |  | 25 | 5.10 | -9 -4 17 |
| Right caudate and thalamus |  | 10 | 4.73 | 12 5 71 |
| Right supramarginal gyrus | BA39 | 23 | 4.99 | 36 -46 35 |
| Right orbitofrontal cortex | BA10/47 |  | 4.90 | 45 20 -1 |

*Note:* Activations are significant at a cluster-corrected FWE of $p < 0.05$ using nonparametric permutations

## Notes

### Competing Interest Statement

The authors have declared no competing interest.

